# Synergistic Effect of Valproate and Constraint Induced Movement Therapy on Functional Recovery Following Ischemic Stroke in Rats: Neuroplasticity and Corticosterone Associated Mechanisms

**DOI:** 10.64898/2025.12.23.695091

**Authors:** P.G. Rini, Yoganarasimha Doreswamy, T.R. Laxmi

**Affiliations:** Department of Neurophysiology, National Institute of Mental Health and Neurosciences (NIMHANS), Institute of National Importance, Bengaluru – 560 029, Karnataka, INDIA

**Keywords:** Ischemic stroke, Sodium Valproate, Constraint Induced Movement Therapy, Neural plasticity, Local field potential recording, Plasma corticosterone, HDAC inhibitors

## Abstract

Most rehabilitation paradigms primarily rely on physical stimuli, but biological enhancers may play a complementary role in optimizing adaptive outcomes. The HDACi (Histone deacetylase inhibitor) drug valproate has shown neuroprotective properties following cerebral ischemia, but how it interacts with rehabilitative training is poorly understood. Therefore, we investigated the synergistic effect of Valproate (VPA) and constraint induced movement therapy (CIMT) on functional recovery following stroke. We induced cerebral ischemia by middle cerebral artery occlusion in adult male Sprague-Dawley rats. VPA (300 mg/kg, subcutaneously) was administered within 3 hours after reperfusion and continued for 7 days, followed by CIMT for another week (6 hours/day). The study revealed a significant increase in total infarction volume in the stroke rats compared to controls on day 7 post-stroke. The behavioral assessments revealed that combined treatment with VPA and CIMT restored the neurological and motor behavior deficits. VPA treatment reduced plasma corticosterone levels and modulated heart rate variability indices in stroke rats, suggesting involvement of neuroendocrine stress responses in post stroke recovery. The electrophysiological findings reveal improved functional connectivity between the penumbral (left) motor cortex and the uninjured motor cortex following combinational treatment. The present study also examined the effects of chronic CIMT and VPA monotherapies and delayed onset CIMT (one week after stroke) on functional recovery following stroke; however, these approaches showed limited efficacy. Thus, the study concludes that VPA, along with rehabilitative training, may represent a novel strategy to improve stroke induced persistent functional deficits.

**Highlights:**

- Valproate may act as an adjuvant biological enhancer for physical rehabilitation following ischemic stroke.
- Combinational therapy, when administered for two weeks following ischemic stroke, resulted in significantly greater behavioral recovery than monotherapies.
- Valproate did not reduce infarct volume, but modulated heart rate variability indices.
- Valproate treatment attenuated the post stroke elevation of plasma corticosterone.
- An increased power spectral density, along with increased waveform synchronization between the injured and the uninjured motor cortices, was observed during fine motor movements following two weeks of combinational treatment.

## 1. Introduction

Ischemic stroke is classified as one of the leading causes of motor disability with limited potential for spontaneous recovery (1,2). Post-stroke recovery heavily relies on the effectiveness of rehabilitation. However, current rehabilitative training emphasizes mechanical or task-oriented training to enhance experience-dependent plasticity. However, persistent functional deficits and several unmet clinical needs in patients emphasize the requirement of combinational approaches harnessing the brain’s intrinsic capacity for reorganization (3–6). Clinically relevant biological enhancers can augment the rehabilitation-induced functional recovery by modulating brain plasticity-associated mechanisms and stress related pathways involved in post stroke recovery.

The HDAC inhibitor drug valproate(VPA) could be useful for treating different conditions, such as bipolar disorder (7,8), migraines (9,10), mood-related disorders (11,12), epilepsy (13,14), and cancer treatment (15,16),motor-related disorders (17). It has been used as an anxiolytic agent (extreme panic attacks without any pathology (18), generalized anxiety disorder (19), bipolar depression (20), panic disorder(21), post-traumatic stress disorder (22), due to its GABA - enhancing action. Studies using *in vitro* and animal models have shown multimodal actions of VPA, including attenuating glutamate excitotoxicity (23), anti-inflammatory responses (24), HDAC inhibition (25), increased acetylation (26), GABA enhancement (27) and reducing microglial activation (28), thus indicating VPA as a neuroprotective agent. VPA’s anti depressive effect is at least partly related to improving the HPA axis and elevating the expression of BDNF (29) Studies have shown that sub chronic administration of VPA causes a reduction in CRF mRNA expression in the paraventricular nucleus (PVN) and the central nucleus of the amygdala (CeA) (30). During pro-apoptotic conditions, either overexpressing CBP (CREB binding protein)(31) or down-regulating HDACs (32,33) is beneficial for neuronal protection at acute time points. In agreement with these observations, HDACi drugs showed neuroprotective effects by attenuating apoptosis in the penumbra area of the infarct (34,35). VPA has also been shown to mediate neuroprotection during traumatic brain injury in vivo, which can be through activation of the Nrf2/ARE pathway, a known signaling pathway for counteracting oxidative stress (36).

Rehabilitative strategies for Upper and lower limb disabilities follow the principles of repetition and experience dependent motor learning. Constraint induced movement therapy is considered one of the most effective physical therapies in clinical settings to improve motor disability (37). Previously, we have shown that constraint-induced movement therapy (CIMT) after one week of MCAO was beneficial in ameliorating ischemic stroke-induced deficits in skilled fine motor movements and associated changes in structural plasticity in the penumbral motor cortex (37). According to literature, CIMT promote contra lesional oriented structural and hemispheric connectivity (38,39). However, the effectiveness of most rehabilitative strategies depends upon the active engagement and participation of patients

Psychological stress is a multidimensional phenomenon; the prevalence of depression like behaviors after a stroke is between 24 and 40%(40) and anxiety affects between 22 and 25% of total stroke survivors(41). As per the literature, stroke negatively affects depression and emotional distress(42,43). Studies showed activation of pathological sympathetic over activity called Paroxysmal sympathetic hyperactivity (PSH)(44), can negatively affect stroke recovery. We largely ignore the effect of psychological stress on stroke rehabilitation. Activation of stress signaling pathways negatively impacts the neuroplasticity and can also initiate additional deleterious signaling pathways.

However, no studies have explored whether valproate enhances CIMT induced recovery by modulation of neuroplasticity associated mechanism and corticosterone signaling following ischemic stroke.

## 2. Materials and Methods

### 2.1. Animals

Male Sprague Dawley rats (360-400g) aged 2-3 months old, procured from the Central Animal Research Facility (CARF), NIMHANS, Bengaluru. They were kept in a standard polypropylene cage (22.5cm×35.5cm×15 cm) at a temperature of 25°C and light-controlled (12h light: 12h dark cycle) conditions. They had access to ad libitum food and water. These rats were handled regularly before subjecting them to stroke surgery. The experimental procedure described in the present study has obtained the Institutional Animal Ethics Committee’s (IAEC) approval (Approval ID: AEC/75/498/NP). All efforts were made to avoid stress and to minimize animal suffering.

### 2.2. Groups

Rats were divided randomly into six groups (n=6/group): (1) Sham control (SC), (2) MCAO followed by 0.9% saline injection from day 0 to day 14 (ISC+VC), (3) MCAO with CIMT from day 1 to day 14 post-stroke (ISC+CIMT), (4) a combination of 0.9% saline injection from day 0 to day 7 post-stroke and CIMT from day 8 to day 14 post-stroke (ISC+VC+CIMT), (5) MCAO with VPA injection from day 0 to day 14 (ISC+VPA), (6) Combinational treatment group was with MCAO followed by VPA treatment from day 0 to day 7 post-stroke and CIMT from day 8 to day 14 post-stroke (ISC+VPA+CIMT).

### 2.3. Middle Cerebral Artery Occlusion (MCAO) and Treatments

The ischemic stroke in rats was induced by occluding the middle cerebral artery (MCA) as described previously (45). In brief, a silicone rubber-coated nylon monofilament (Doccol Corporation, USA, diameter with coating 0.41+/-0.02 mm, length 30mm) was inserted through an incision in the internal carotid artery (ICA) and advanced toward the cranial base until slight resistance was felt (approximately 23 ± 2 mm from the bifurcation of the common carotid artery (CCA). The MCAO lasted for 60 minutes, after which the filament was removed for the reperfusion. The SC rats were subjected to vascular surgery without occluding the MCA.

Sodium Valproate (300 mg/kg, s.c, twice daily(46) administered as previously described (Sigma Aldrich, USA). VPA, dissolved in 0.9% normal saline, while the vehicle used was 0.9% saline (w/v) (Abaris Healthcare Pvt.Ltd, India). CIMT was administered for 6h (16.00h – 22.00h) daily (37). Both sodium Valproate and 0.9% saline were administered subcutaneously twice daily to their respective groups, starting within 3 hours after MCAO.

### 2.4 Assessment of Heart Rate Variability (HRV) in Conscious Rats

The functional status of the autonomic nervous system was carried out in rats by measuring the HRV assessment on day 8 of post-stroke surgery for a period of 15 minutes. The ECG needle electrodes were inserted subcutaneously in the right and left forelimbs, and into the left hind limb (Lead Ⅱ configuration). ECG signals acquired from the amplifier were digitized at a sampling rate of 1000 Hz and analyzed (Lab chart 8 software, AD Instruments, USA) from 5 minutes’ epoch. The mean HRV, LF, HF, LF/HF, and total power were calculated through time domain and frequency domain analysis.

### 2.5. Plasma Corticosterone

Plasma corticosterone was assessed after 7 days of MCAO using the ELISA kit method (PuregeneTM, Genetix Biotech Asia Pvt. Ltd, #PG-9430 M) (47)

### 2.6. The 2, 3, 5 Triphenyltetrazolium Chloride Staining (TTC)

The rat brains after 7 days of stroke were placed in a brain matrix slicer to obtain 2 mm coronal sections and immediately exposed to TTC staining to measure infarct volume in the injured cortex. The infarct volume was measured as described previously (37).

### 2.7. Stereotaxic Surgery and Local Field Potentials (LFP) Recording During Reach to Grasp Task

The coated stainless steel electrodes (125 μm) (A-M Systems, Inc., USA) were implanted stereotaxically (Stoelting Co., USA) under anesthetized conditions (Ketamine 80 mg/kg + Xylazine 10 mg/kg ) (48) in layer V of the right and the left motor cortex (anterior-posterior (AP), +3.0 mm; mediolateral (ML), ±2.0 mm; and dorsoventral (DV), 1.5 mm). The ground and reference screws were placed subdurally in the right cerebellum and in the left cerebellum, respectively. The MCAO surgery was performed after implantation of electrodes for electrophysiological recordings. Rats were allowed to recover from the surgery for up to 7 days. The electrode placement and brain infarct were confirmed on day 16 post-stroke using the Nissl staining method after transcardial perfusion of the rat brain. The local field potentials (LFP) were recorded during the reach to grasp task on days 8 and 15 post stroke and analyzed using spike 2 (version 5.0) (CED, UK) software system.

### 2.8. Constraint - Induced Movement Therapy (CIMT)

The CIMT was administered as described in our previous study (37). Briefly, the cast was applied manually around the uninjured forelimb. The cast was prepared using plaster of Paris. All animals that received CIMT therapy were subjected to reach to grasp task training, promoting the use of the injured limb.

### 2.9. Neurological Examination

Neurological examination was conducted one day before, on day 8, and on day 15 post-MCAO, using the Garcia scoring method. The scoring method was done as per the previous studies. (49).

### 2.10. Cylinder Test

A cylinder test was used to assess the spontaneous forelimb paw preference in a natural environment (50) and was conducted on days 8 and 15 post-MCAO in rats. The rat was placed in a Plexiglass cylinder (34cm x 21cm x 40cm) and allowed to explore the cylinder wall for 5 minutes, during which video recordings were made for the offline analysis. The limb preference was measured as the ipsilateral bias index and calculated using the following formula:

Ipsilateral bias index = i-c /i+ b+c

Where i, represents the number of independent unimpaired forelimb contacts; c is the number of dependent impaired forelimb contacts; b, is the number of both forelimbs in contact simultaneously.

### 2.11. Rotarod Test

The Rotarod test was carried out to assess the balance and coordination in rats by using the Rotamex-5 system (CR-1 Rotamex System, Columbus Instruments, USA). All the rats were subjected to 5 days of training before the stereotaxic surgery and continued 5 days after MCAO upto day 14 of post-MCAO. The test was carried out on days 8 and 15 post stroke.The maximum speed at which the animal stayed for a given duration was measured through Rotamex software. The longest duration of stay in the rotating rung was considered a measure of balance and coordination in the rotating wheel (51).

### 2.12. Gait Analysis

Rats were allowed to walk in a narrow passage on plain paper one meter in length. An ink was applied to the ventral side of the forepaw to get a footprint. Then, stride length was measured by calculating the distance between two consecutive footprints on the same side.

### 2.13. Reach to Grasp Task

The Reach to grasp task was used to assess and train skilled fine forelimb movements. The behavioural procedure was carried out as described previously (37). Briefly, the rats received two weeks of training, with 30 pellets for 30 minutes (1 pellet/minute). Rats that learned the reach to grasp task were selected for stereotaxic surgery, followed by MCAO surgery. The training continued once again after the MCAO surgery for 2 weeks without food deprivation. The test was performed on days 8 and 15 post stroke for 10 minutes (1 pellet/minute). The videos were recorded and analyzed offline using a VLC media player. The percentage success was calculated for each day.

## 3. Statistical analysis

All data is expressed as mean ± SEM. One-way ANOVA followed by multiple comparison test was used to assess the valproate effect on infarct size, plasma corticosterone, and heart rate variability parameters. To check the effect of different therapeutic strategies (monotherapy vs combinational therapy) on functional recovery after stroke, a repeated measures ANOVA followed by Tukey’s multiple comparison test was performed. The data were analyzed using GraphPad Prism software (version 9). The criterion for statistical significance was at a probability level of p<0.05.

## 4. Results

VPA, in conjunction with CIMT, promoted functional recovery, showing an increased neuronal connectivity between the injured penumbral and uninjured motor cortex during the reach-to-grasp task.

### 4.1. Infarct Size, Cardiac Functions

Figure 2a shows the TTC-stained images of sham control (SC) and vehicle control (ISC+VC) and VPA-treated (ISC+VPA) rat brain sections after 7 days following MCAO. The affected area in the stroke brain appears as white after TTC staining. One-way ANOVA followed by Tukey’s multiple comparison test revealed an increased infarct volume in the ISC +VC (p<0.0001) and ISC+VPA (p=0.0001) rats when compared to SC rats (F_2, 15_=33.15, p<0.0001) (Figure 2b). No significant difference in infarct volume was observed in the ISC+VPA group compared with the ISC+VC group.

**Figure 1:**
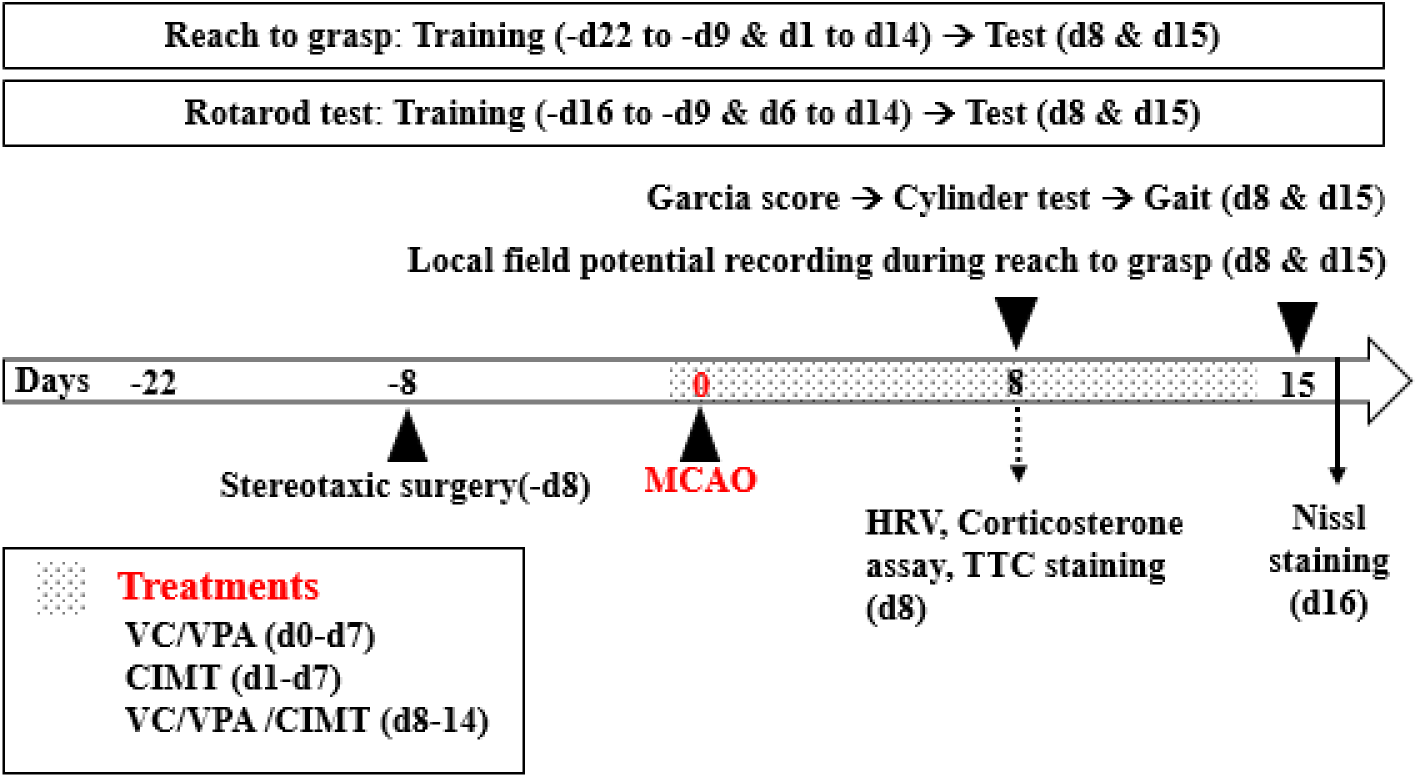
A study design depicting the timeline of the procedures carried out

**Figure 2:**
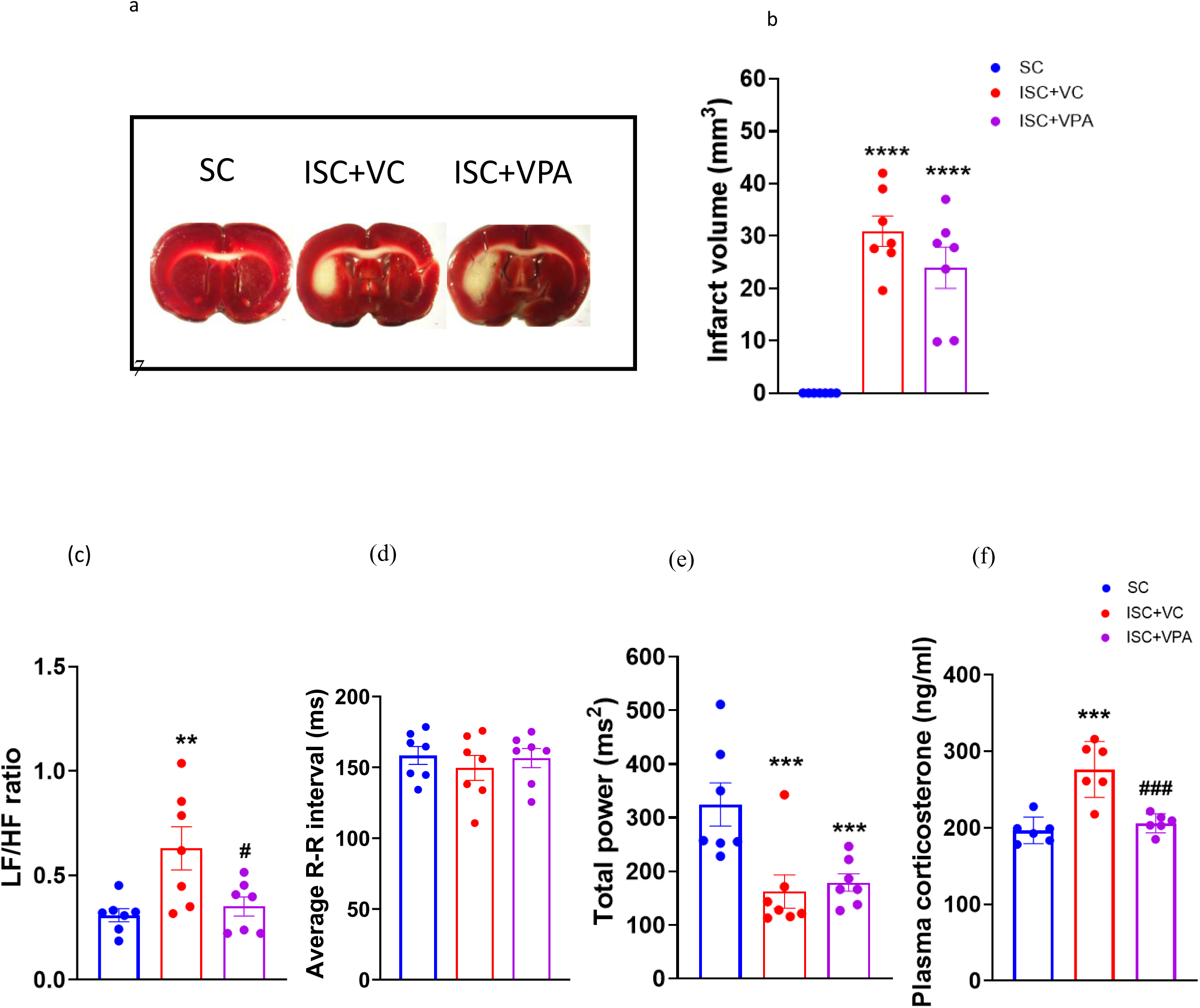
The ischemic injury and associated changes in cardiac functions and systemic stress response in the rat after 7 days of MCAO surgery. (a) Representative images of rat brain section showing the infarct size/area, (b) Infarct volume (mm^3^); Cardiac functions showing changes in (c) LF/HF ratio to indicate sympathovagal balance, (d) R-R intervals, (e) Total power, (f) Plasma corticosterone level on day 8 post-stroke from SC, ISC+VC and ISC+VPA rat groups. Data expressed as Mean±SEM. One-way ANOVA followed by a post hoc test showed significant differences between groups. **p<0.01, ***p<0.001, ****p<0.0001 vs SC groups, ^#^p<0.05 ^###^p<0.001 vs ISC+VC (n=6-7/group).

Similarly, the associated changes in cardiac functions on day 8 post-stroke were evaluated using one-way ANOVA followed by post hoc analysis. The results revealed significantly increased LF/HF ratio (F_2,15_=1.4, p<0.01) (Figure 2c), decreased R-R intervals (F_2,15_=.94, p<0.05) (Figure 2d), and total power (F_2,15_=.27, p<0.01) (Figure 2e) in ISC+VC groups when compared to SC groups. In addition, ISC+VPA rats showed a significantly reduced total power (p<0.01) when compared to the SC groups. However, compared to the ISC+VC group, the ISC+VPA groups showed significantly reduced LF/HF ratio(p=0239). Together, these results demonstrated that VPA did not reduce infarct volume but significantly reduced LF/HF ratio.

### 4.2. Plasma Corticosterone

To examine the role of stress on stroke recovery, the plasma corticosterone levels were estimated. There were significantly increased plasma corticosterone levels in ISC+VC compared to SC groups (F_2,15_=18.92, p<0.001), while significantly reduced levels in the ISC+VPA group when compared to ISC+VC (p<0.001) on day 8 post stroke (Figure 2f).

### 4.3. Neurological Score

As shown in Figure 3a, all ISC and rats treated with VPA and/or CIMT groups showed an overall deficit in Garcia score compared to the SC group after one and two weeks of MCAO. The two-way ANOVA with repeated measures of analysis revealed a statistically significant effect on groups (F_5,36_=199.8, p<0.0001) and groups x days (F_5,36_=19.35, p<0.0001). Tukey’s multiple comparison test has further revealed a significant effect on neurological functions, showing significantly reduced Garcia score in ISC+VC (p<0.0001), ISC+CIMT (p<0.0001), ISC+VC+CIMT (p<0.0001), ISC+VPA (p<0.0001), and ISC+VPA+CIMT (p<0.0001) when compared to SC groups on both day 8 and day 15 post-stroke. Compared with ISC+VC, the ISC+VC+CIMT (p<0.01), ISC+VPA (p<0.05) and ISC+VPA+CIMT (p<0.05) groups showed significant improvement on day 15 post stroke.

**Figure 3:**
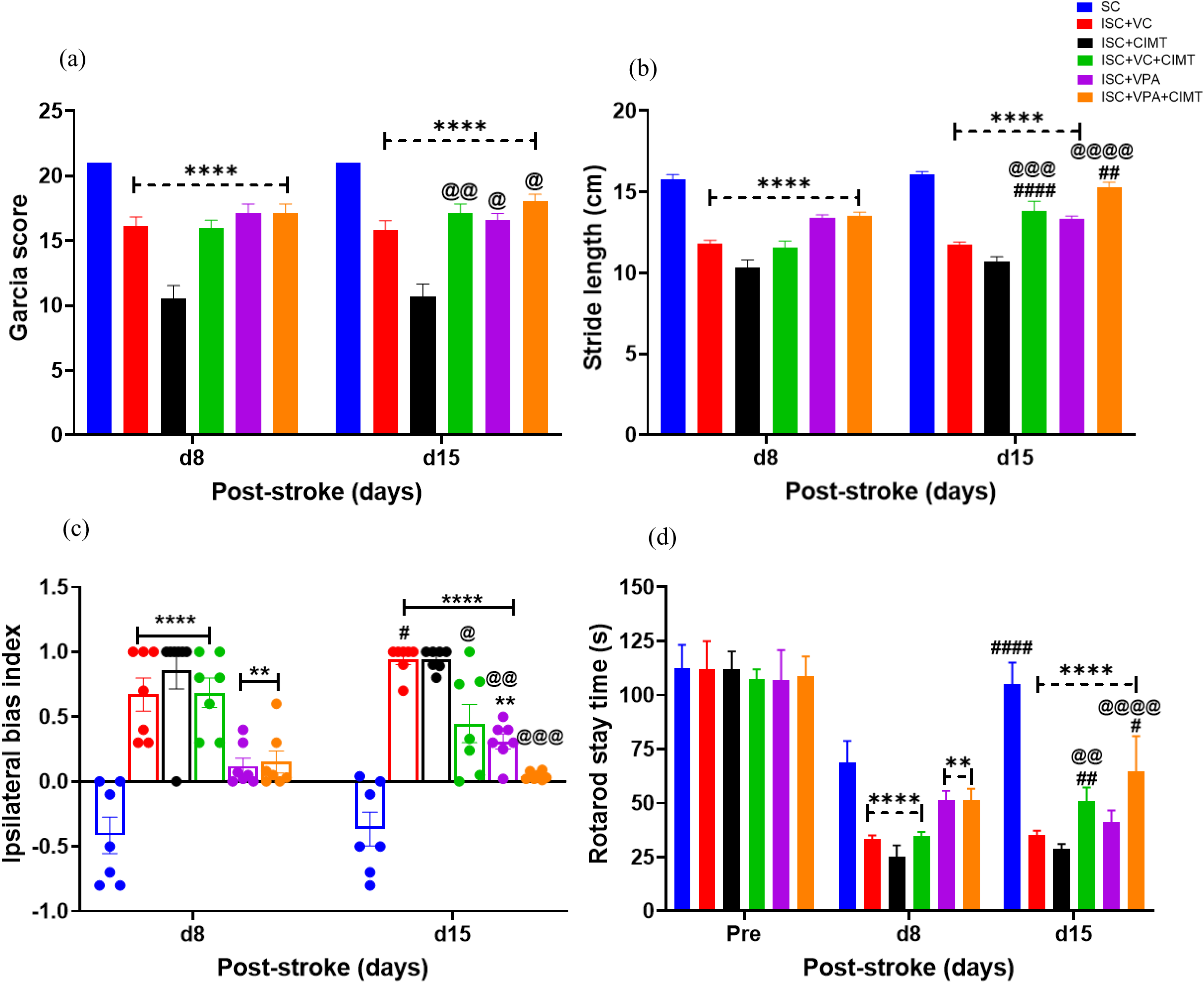
Behavioral assessment on day 8 and day 15 after MCAO surgery. (a) Garcia score, (b) Gait analysis by measuring stride length, (c) Ipsilateral bias index to assess the forelimb preference, (d) Rotarod stay time for gross motor movement assessment – pre and post-MCAO surgery. Data is represented as Mean ± SEM. Two-way ANOVA, followed by Sidak’s multiple comparison test, was performed (n=7/group). ****p<0.0001, **p<0.01 vs SC, ^@@@@^p<0.0001, ^@@^p<0.01, ^@^p<0.05 vs ISC+VC, ^####^p<0.0001, ^##^p<0.01, ^#^p<0.05 vs between day 8 and day 15.

### 4.2. Gait

The stride length was assessed in rats subjected to ischemic stroke for the gait analysis. The two-way ANOVA with repeated measures of analysis revealed a statistically significant effect on groups (F_5,36_=53.07, p<0.0001) and groups x days (F_5,36_=6.494, p<0.001). Sidak’s multiple comparison test has further revealed a significant effect on stride length, showing reduced stride length in ISC+VC (p<0.0001), ISC+CIMT (p<0.0001), ISC+VC+CIMT (p<0.0001), and ISC+VPA (p<0.0001) when compared to SC groups on days 8 and 15(Figure 3b). The ISC+VPA+CIMT group, on the other hand, showed a reduced stride length on day 8 (p<0.0001) but not on day 15 when compared to the SC group. Comparison between post-stroke days 8 and 15 revealed a significant improvement in stride length in both the ISC+VC+CIMT (p<0.0001) and ISC+VPA+CIMT (p<0.0001) groups, but not with the other groups. The ISC+VC+CIMT (p<0.0001) and ISC+VPA+CIMT (p<0.0001) groups exhibited a significantly greater stride length on day 15 post-stroke compared with the ISC+VC group (Figure 3b).

### 4.4. Forelimb Use

The forelimb use asymmetry in rats subjected to ischemic stroke was assessed using a cylinder test. The two-way ANOVA with repeated measures of analysis revealed a statistically significant effect on groups (F_5,36_=34.25, p<0.0001) and groups x days (F_5,36_=2.568, p<0.05). Sidak’s multiple comparison test has further revealed a significant effect on the ipsilateral bias index showing increased ipsilateral bias index score in ISC+VC (p<0.0001), ISC+CIMT (p<0.0001), ISC+VC+CIMT (p<0.0001), and ISC+VPA (p<0.01) when compared to SC groups on day 8. The ISC+VPA+CIMT group, on the other hand, showed an increased ipsilateral bias index on day 8 (p<0.01) but not on day 15 when compared to the SC group. When comparisons were made between post-stroke days on 8 and 15, there were significant differences found in ISC+VC (p<0.05), but not with other groups (Figure 3c).

### 4.5. Gross Motor Movements

The rotarod test was performed at baseline and on post-stroke days 8 and 15. The two-way ANOVA with repeated measures of the analysis revealed a statistically significant effect on groups (F_5,36_=50.34, p<0.0001) and groups x days (F_10,72_=24.06, p<0.001). Tukey’s multiple comparison test has further revealed a significantly reduced rotarod stay time on day 8 and day 15 in ISC+VC (p<0.0001), ISC+CIMT (p<0.0001), ISC+VC+CIMT (p<0.0001), ISC+VPA (p<0.01), and ISC+VPA+CIMT (p<0.01) when compared to SC groups. When comparisons were made between post-stroke days 8 and 15, we found significant differences in rotarod stay time in SC (p<0.0001), ISC+VC+CIMT (p<0.01), ISC+VPA+CIMT (p<0.05), but with no differences in ISC+VC, ISC+VPA and ISC+CIMT groups. Compared with the ISC+VC group, the ISC+VC+CIMT (p<0.01) and ISC+VPA+CIMT (p<0.0001) groups showed significant improvement on day 15 post-stroke (Figure 3d).

### 4.6. Fine Skilled Movements

The two-way ANOVA with repeated measures of analysis revealed a statistically significant effect on groups (F_5,36_=197.8, p<0.0001) and groups x days (F_5,36_=6.655, p<0.001). Tukey’s multiple comparison tests have further revealed a significantly reduced reach success rate on day 8 and on day 15 in ISC+VC (p<0.0001), ISC+CIMT (p<0.0001), ISC+VC+CIMT (p<0.0001), ISC+VPA (p<0.0001), and ISC+VPA+CIMT (p<0.0001) compared to SC groups. When compared to day 8, the reach success rate on day 15 was significantly increased in ISC+CIMT (p<0.01), ISC+VC+CIMT (p<0.001), ISC+VPA (p<0.05), and ISC+VPA+CIMT (p<0.0001), but with no differences in ISC+VC and SC groups. Additionally, we observed a significant improvement in skilled reaching in the ISC+CIMT (application of CIMT immediately after stroke) from day 8 to day 15 post stroke (p<0.01). This indicates that chronic treatment for two weeks with VPA and/or CIMT was beneficial in restoring skilled fine motor movements in ischemic stroke rats (Figure 4).

**Figure 4:**
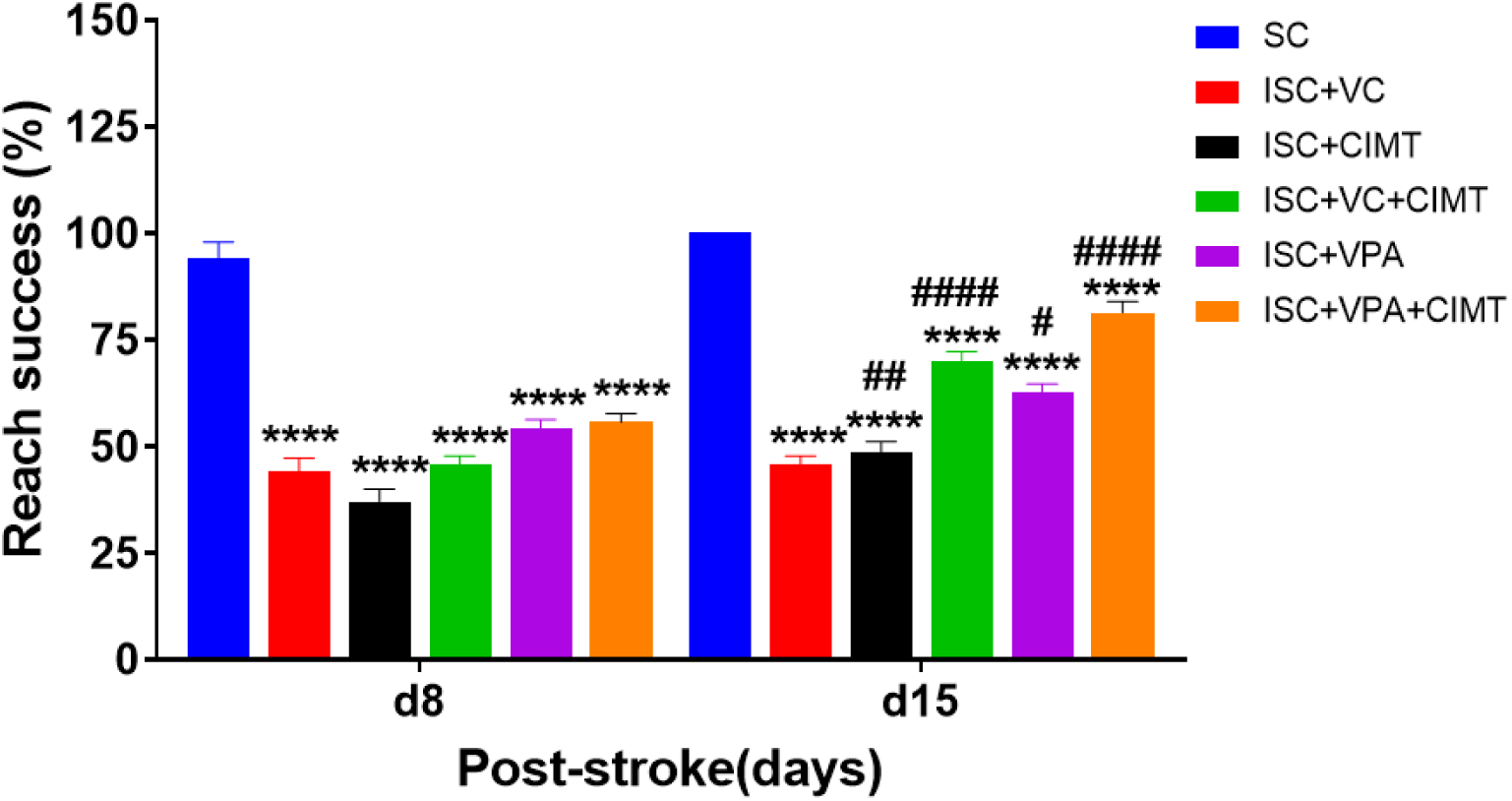
Reach to grasp task to assess the skilled forelimb movements. Reach success rate on day 8 and on day 15 post-MCAO surgery. Data is represented as Mean ± SEM. Two-way ANOVA followed by Tukey’s multiple comparison test (n=7/group). ****p<0.0001 vs SC, ^####^p<0.0001, ^##^p<0.01, ^#^p<0.05 between day 8 and day 15.

**Figure 5:**
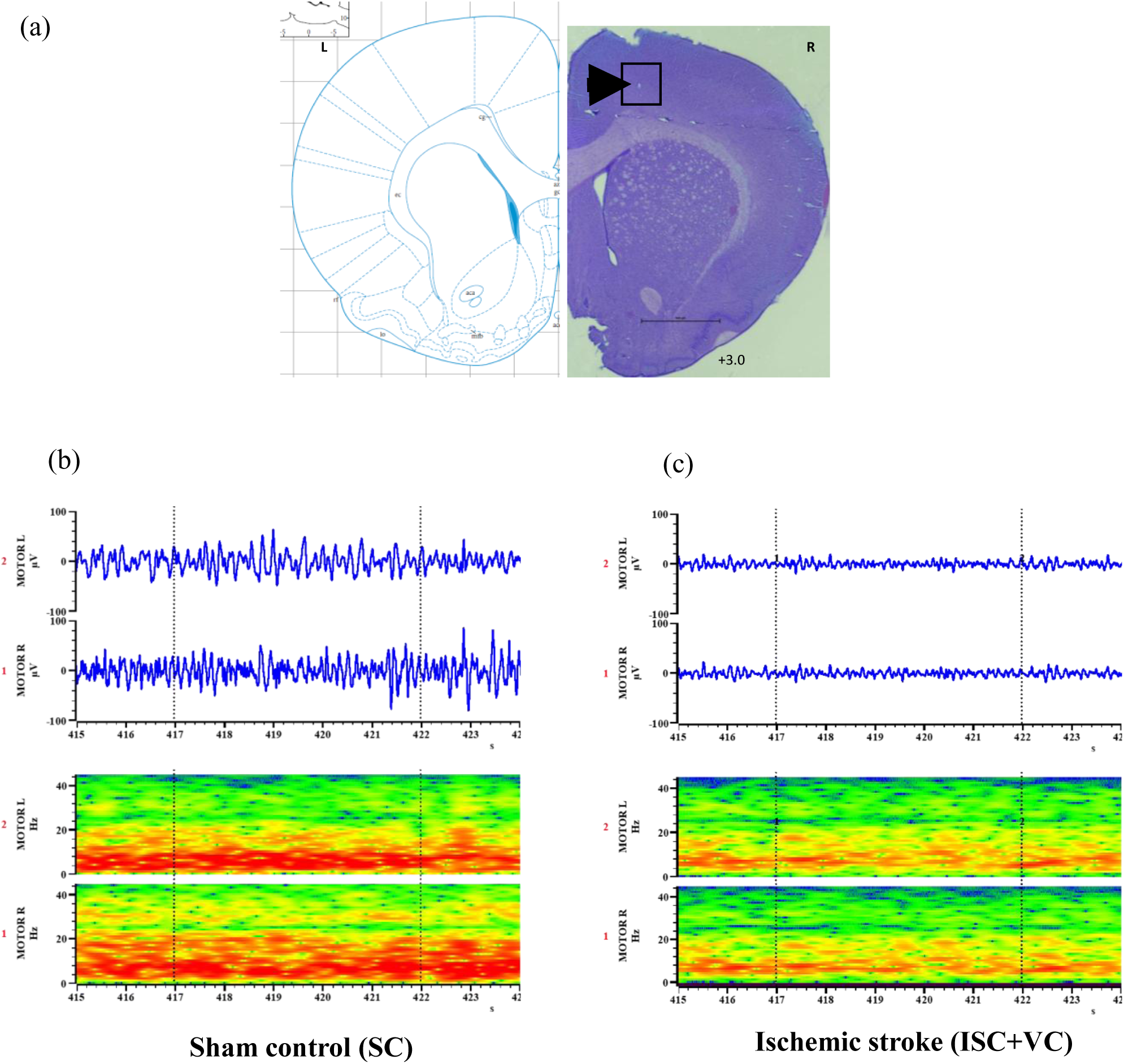
Histological verification of electrode placement in rat motor cortex. (a) Confirmation of electrode placement for local field potential (LFP) recording showing electrodes in layer 5 of the motor cortex. (b) Representative examples of LFP recordings and spectrogram from right and left motor cortex during the skilled forelimb movement from (b) Sham control (SC) and (c) Ischemic stroke (ISC+VC) rat groups. The event marker shows the event selected to analyze the LFPs during the specific phase of the reach to grasp task.

### 4.7. Electrophysiological Correlates of Fine Skilled Movements

#### 4.7.1. Power Spectral Analysis (PSD)

Oscillatory activity in the motor cortex was classified into the following frequency bands: Delta (0–4 Hz), Low theta (4–8 Hz), High theta (8–12 Hz), Beta (12–20 Hz), and Gamma (20–40 Hz) frequency bands. The statistical analysis was carried out using two-way ANOVA with repeated measures of analysis followed by Tukey’s multiple comparison tests.

At the delta frequency band, there were no significant differences in relative power between groups and days in both the injured (Figure 6a) and the uninjured motor cortex (Figure 6e).

**Figure 6:**
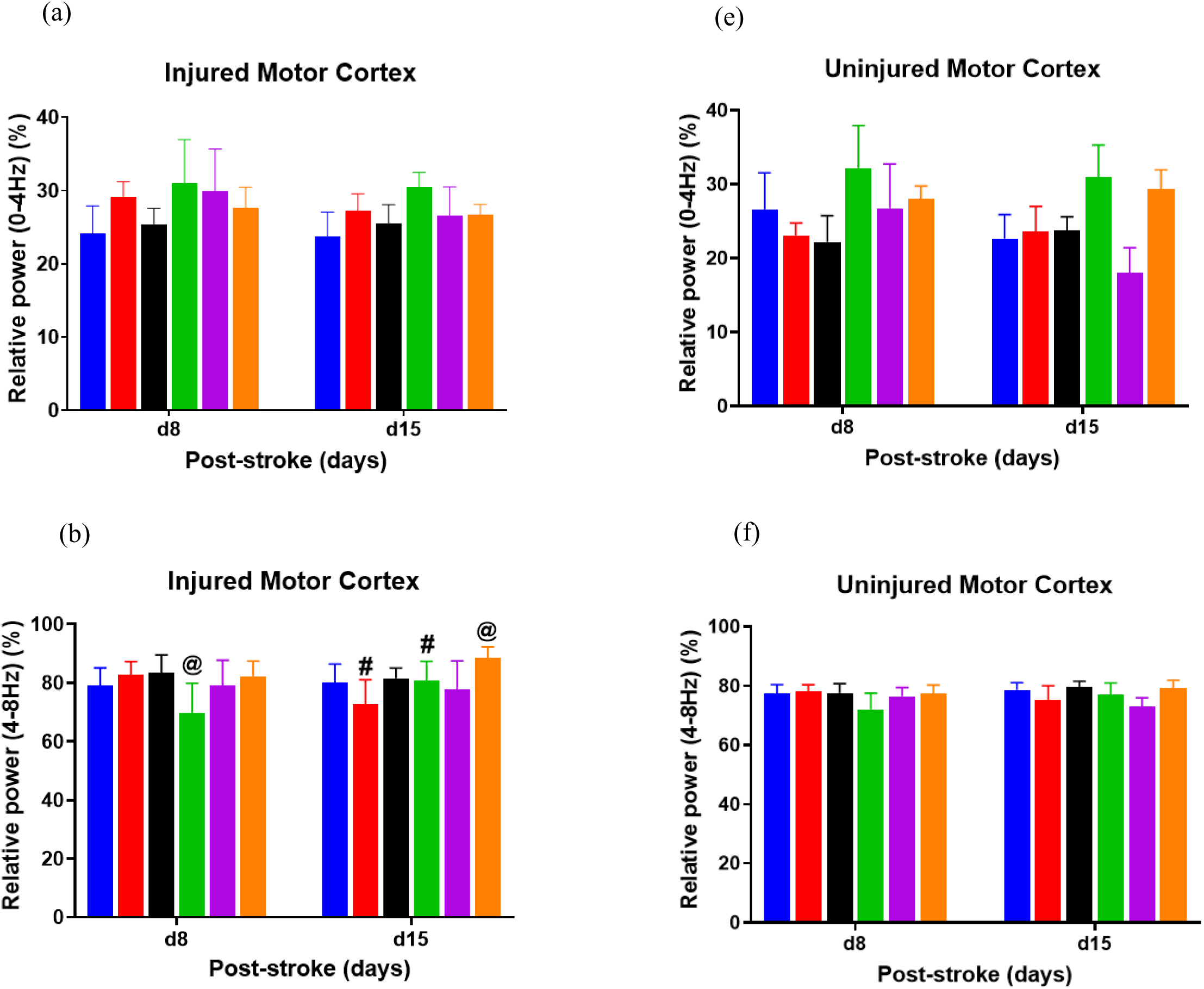
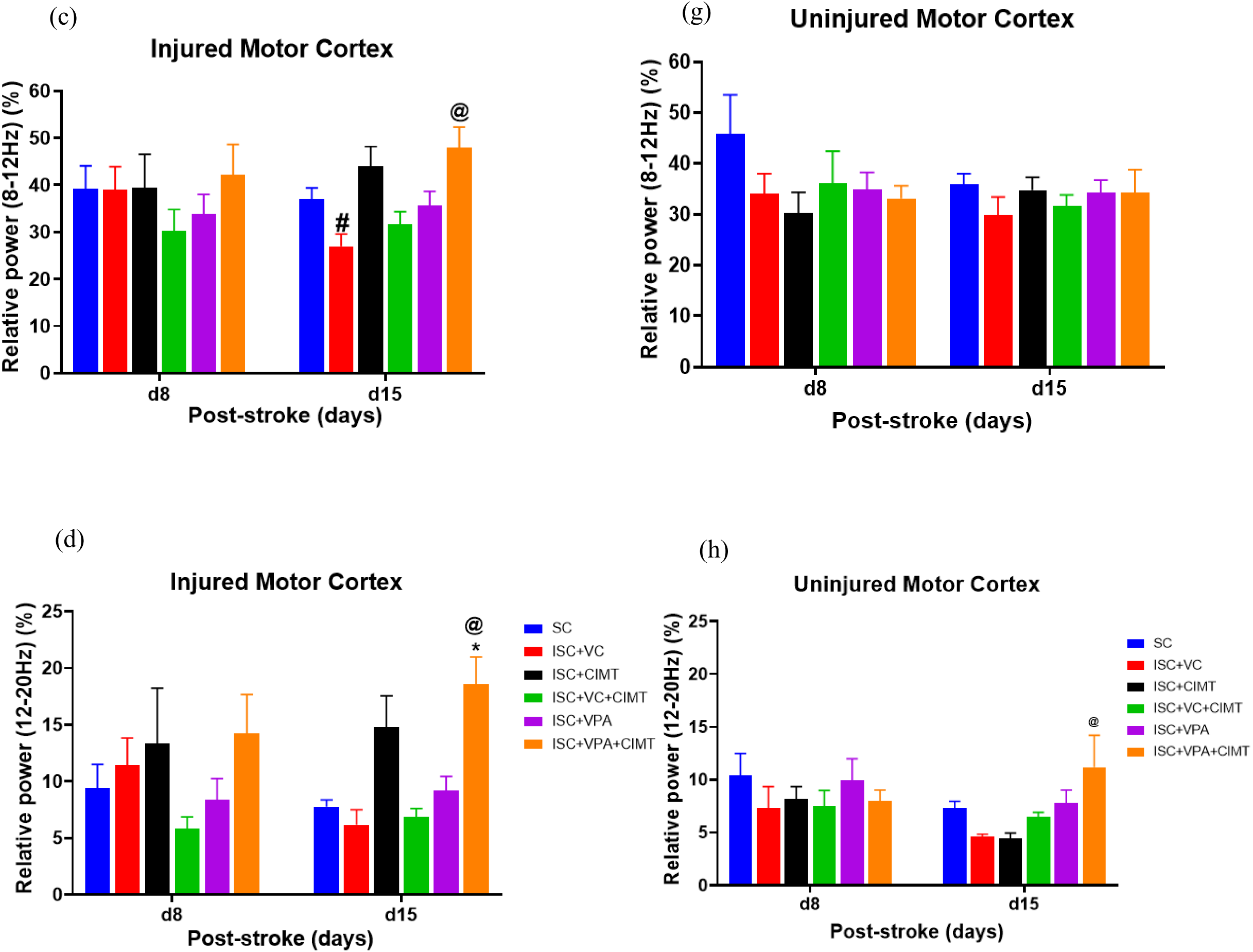
Averaged Power Spectral Density (PSD) on day 8 and day 15 following MCAO from the injured (left) and the uninjured (right) motor cortex. Relative Power (%) at (a) 0-4Hz, (b) 4-8 Hz, (c) 8-12Hz, and (d) 12-20 Hz frequency band in the injured motor cortex; (e) 0-4Hz, (f) 4-8Hz, (g) 8-12Hz and (h) 12-20Hz in the uninjured motor cortex following 7 and 14 days of treatment with VPA+/CIMT. Two ways ANOVA followed by Tukey’s multiple comparison tests was performed. Data is represented as Mean±SEM. *p<0.01 vs SC, ^@^p<0.05 vs ISC+VC and ^#^p<0.05 between day 8 and day 15 post-stroke, (n=6/group).

The two-way ANOVA with repeated measures on theta frequency band (4-8Hz) has revealed a significant effect on group (F_5,30_=2.802, p<0.05) and post-stroke days versus group (F_5,30_=3.9476, p<0.01) on day 8 post-stroke. Tukey’s multiple comparison test has further revealed a significantly increased theta power at 4-8Hz in ISC+VPA+CIMT (p<0.05) compared to the ISC+VC+CIMT group on day 8 post stroke. There was also a significant increase in theta power on day 15 post-stroke in ISC+VPA+CIMT compared to ISC+VC groups (p<0.003). The significant difference was found between day 8 and day 15 post-stroke in ISC+VC and ISC+VC+CIMT rat groups (Figure 6b). However, there were no differences observed in the uninjured motor cortex (Figure 6f).

Tukey’s multiple comparison test has revealed that the significant differences in PSD was observed at high theta frequency band (8-12 Hz) on day 15 between ISC+VC and ISC+VPA+CIMT in the injured motor cortex (p<0.05) (Figure 6c). Similarly, between day 8 and day 15, there were significant differences in the ISC+VC group (p<0.05) at the high theta band. The significant differences were not observed in the uninjured motor cortex (Figure 6g).

In the injured motor cortex, the two-way ANOVA was carried out to identify the impact of treatment with VPA and CIMT at the beta frequency band (12-20Hz), which revealed a significant effect on groups (F_5,30_=4.198, p<0.01). Tukey’s multiple comparison test has further revealed that the PSD was significantly increased in ISC+VPA+CIMT when compared to SC (p<0.01), ISC+VC (p<0.01), and ISC+VC+CIMT (p<0.01) on post-stroke day 15, but not on day 8 in the injured motor cortex (Figure 6d). Similar observations were observed in uninjured motor cortex (Figure 6h), wherein the ISC+VPA+CIMT group showed significantly increased PSD at 12-20 Hz (p<0.05) on day 15 but not on day 8 when compared to ISC+VC and ISC+CIMT.

Next, the power spectral density (PSD) analysis of the left and motor cortices was carried out based on the area under the curve (AUC) during the reach to grasp task for day 8 and day 15 post-stroke in rats. The analysis with two-way ANOVA has revealed a statistically significant effect on groups (F_5,30_=5.016, p<0.001). Sidak’s multiple comparison test has further revealed significantly increased PSD in the injured (left) motor cortex during reach to grasp task (Figure 7a) on day 15 in ISC+VPA+CIMT when compared to SC (p<0.05), ISC+VC (p<0.001), ISC+VC+CIMT (p<0.05), and ISC+VPA (p<0.05) groups. When compared to day 8, there were significant differences (p<0.05) on day 15 found in the ISC+VC rat groups. In the uninjured (right) motor cortex, on the other hand, we did not find any significant differences in PSD between groups and post-stroke days (Figure 7b).

**Figure 7:**
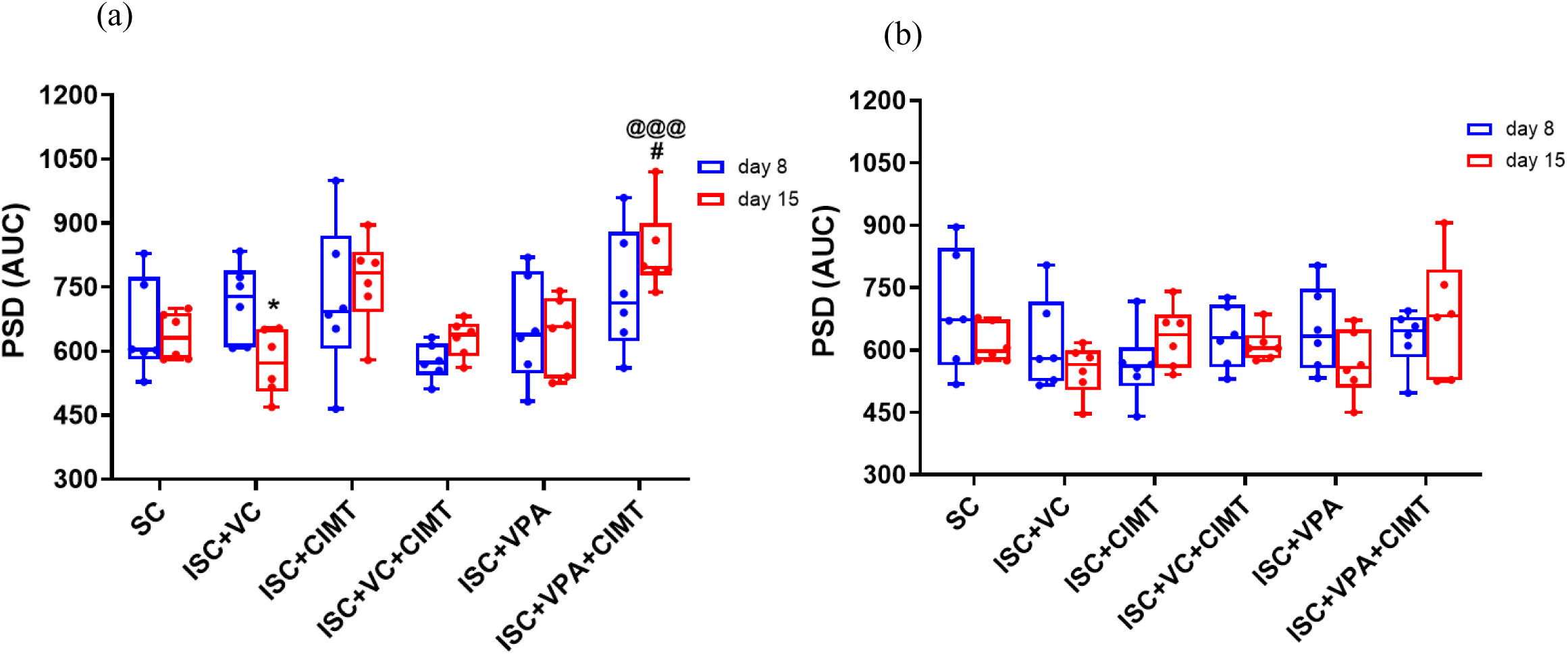
Averaged PSD based on area under the curve (AUC) on day 8 and day 15 following MCAO from the (a) injured (left) and (b) the uninjured (right) motor cortex after 7-14 days of treatment with VPA+/CIMT. Two-way ANOVA followed by Tukey’s multiple comparison test. Data is represented as Mean±SEM. *p<0.05 vs SC; ^#^p<0.05, PSD (AUC) between d8 and d15, ^@@@^p<0.001, PSD (AUC) as compared to ISC+VC.

#### 4.7.2. Interhemispheric Connectivity

To investigate the effects of combinational therapy on interhemispheric functional connectivity, the waveform cross-correlation analysis was carried out on LFP data for the day 8 and day 15 post-stroke rehabilitation periods. As compared to the SC group, there was a significant effect on the group (F_5,30_ =11.01, p<0.0001) on day 15 and not on day 8 post-stroke period. Sidak’s multiple comparison test has further revealed that the peak cross-correlation was significantly reduced more in ISC+VC (p<0.0001), ISC+CIMT (p<0.0001), ISC+VC+CIMT (p<0.01), ISC+VPA (p<0.01), and less significantly in ISC+VPA+CIMT (p<0.05) when compared to the SC group of rats. On the other hand, the significant differences in peak correlation level were not observed on day 8 post-stroke. The peak correlation between day 8 and day 15 was significantly greater in the SC rat group (p<0.05) and not observed in other stroke groups. Thus, these data together indicate that the synchronization between hemispheres was gradually built as the rat acquired skilled fine forelimb motor movements. Similarly, the cortical synchronization level was impacted on day 15 of the post-stroke treatment strategy, suggesting an increased association between interhemispheric connectivity of an increased reach to grasp task (Figure 8).

**Figure 8:**
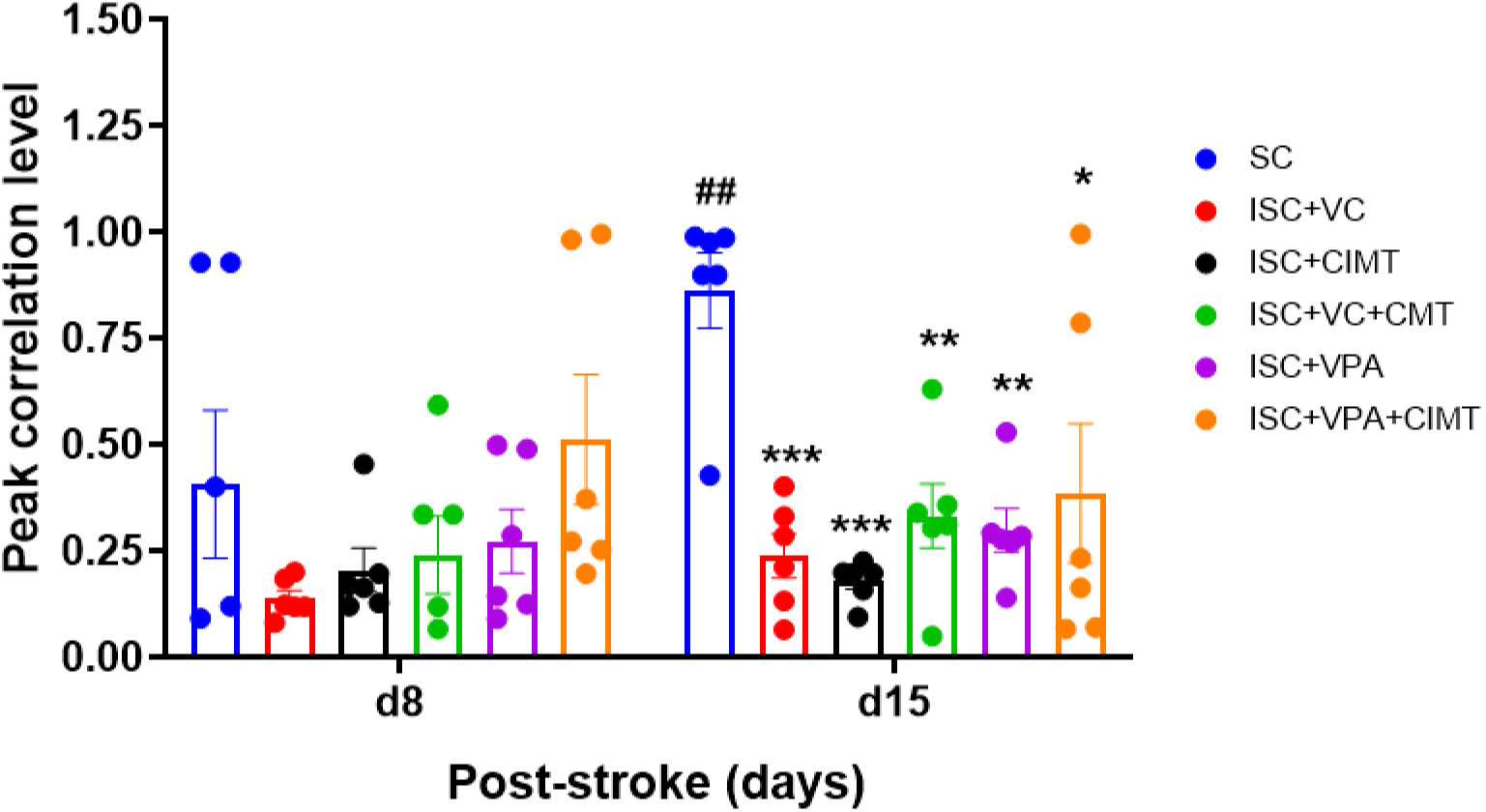
Waveform correlation analysis. Peak correlation level indicates the synchronization level between the left and the right motor cortex. Data expressed as Mean±SEM, Two-way ANOVA (F_5,30_ = 11.01, p<0.0001) followed by Sidak’s multiple comparison test (n=6/group). Data expressed as Mean±SEM. ***p<0.001, **p<0.01, *p<0.05 vs SC; ^##^p<0.01 between day 8 and day 15

## 5. Discussion

The present study provides behavioral and electrophysiological evidence that early treatment with VPA, combined with the rehabilitative strategy, helps to enhance the functional connectivity between the penumbral cortex and uninjured motor cortex following stroke.

We observed that early administration of VPA after stroke significantly improved the stroke-induced deficits in neurological functions, gait, ipsilateral bias, gross motor movements, and skilled forelimb movements, even though acute VPA treatment did not reduce infarct volume. These improvements are further enhanced when combined with rehabilitation. An increased LF/HF ratio and reduced total power, suggesting a reduced cardiovascular fitness in vehicle control rats, when compared to sham control rats, following ischemic stroke. However, one week of Valproate treatment significantly improved sympathovagal balance compared to the vehicle control group. The reduction in plasma corticosterone levels following Valproate treatment suggests a role for stress hormone modulation in stroke recovery. Taken together, VPA was beneficial in reducing neuroendocrine stress response following experimental stroke.

### 5.1. Behavioral Recovery in Response to Combinational Treatment with VPA and CIMT

The neuroprotective effect of HDACi is mainly due to its role in epigenetic modulation of gene expressions (52), which has enabled the present study to administer the VPA starting 3 h after MCAO and continued for 7-14 days to treat ischemic stroke conditions. The behavioral response to VPA is possibly mediated by inhibiting the inflammatory and oxidative pathways (25). It was shown previously that the delayed application of CIMT for a week has ameliorated the stroke-induced deficits in fine motor movements and increased the quality of movements by increasing dendritic arborization in the pyramidal neurons of layer III and layer V motor cortex (37). According to the literature, immediate application of CIMT following ischemic stroke can have a detrimental effect on the functional recovery (53). The present study has substantiated these findings, showing reduced behavioral recovery in rats subjected to CIMT within 3 hours after MCAO, indicating that early physical rehabilitative interventions may be more harmful to the ischemic stroke recovery process. On the contrary, the delayed application of CIMT was beneficial in restoring the behavioral deficits induced by MCAO, which is in line with our previous findings (37).

Neuroprotective monotherapies enhance functional recovery in stroke-induced rats (54). Comparison between the two monotherapies revealed that early VPA treatment was more effective in reducing ipsilateral bias than early CIMT therapy. The reinforcing effect of late CIMT combined with VPA significantly improved gross motor and gait deficits post-MCAO, highlighting the importance of integrating adjuvant therapies along with rehabilitation in stroke treatment.

### 5.2. Combinational Treatment Restores Connectivity Between the Penumbral and Uninjured Motor Cortex During Fine Motor Moments

The functional recovery after stroke requires an anatomical connectivity between the penumbral and uninjured motor cortex (55) during motor functions to predict the neurological and motor functional outcome (56). The combinational treatment after stroke restored stroke-induced reduction in the power spectral density at theta band in the injured motor cortex to normal by day 15 compared to monotherapies during reach to grasp task. The benefits of combinational treatment extended to an increased waveform synchronization between the right and left motor cortices from day 8 to day 15 after the onset of ischemic stroke indicating the functional recovery in fine forelimb movements.

The fine motor movements in rodents are primarily controlled by neural circuits involving the motor cortex (57). The fine motor skills are typically the most delayed motor functions to recover from the stroke (58). In several cases, even after rehabilitation therapies, patients are left with persistent fine motor deficits (59). It is interesting to note that stroke-induced deficits in the skilled fine motor movements is long-lasting, but not permanent due to direct association with the neural plasticity in the penumbral motor cortex and associated with the restoration of interhemispheric connectivity. This suggests that neuroplasticity adaptations may require much more time before translating into measurable, true motor recovery measures.

Enhanced synchronization at theta and beta frequency range during reach to grasp task indicates that low-frequency oscillations such as theta rhythms are critical for long-range communication between motor circuits, compensatory network reorganization (60) indicative of ongoing plasticity and recovery mechanisms (64). In stroke-induced rats, a combinational treatment resulted in increased theta oscillations in the injured M1 (motor cortex) region during the task, reflecting attentional processing of movement control and possibly indicating ongoing reorganization within the penumbra of the infarct. Additionally, an increase in beta power in the injured M1 suggests an enhanced cortical excitability during motor output. However, few studies indicated that increased beta frequency range may suggest maladaptive plasticity (65) with engagement of compensatory cognitive mechanisms during the task (66). The reappearance of these oscillatory patterns with combined treatment, thus, is associated with gradual improvements in fine motor performance (62). These findings support the idea that combinational therapies facilitate neural plasticity in the peri-lesioned areas necessary for fine motor recovery, more likely by directing the neuroprotective qualities of VPA to specific goal-oriented synaptic remodeling and connectivity with the help of CIMT.

Taken together, the results from the present study indicate a strong link between the recovery of fine forelimb movements and frequency-specific cortical reorganization within and between the hemispheres, showing the restoration of interhemispheric connectivity, especially long-range connections, after two weeks of treatment for ischemic stroke. Thus, the study revealed that immediate treatment for ischemic stroke with sodium valproate in conjunction with CIMT appears to enhance the recovery process by reducing the dominance of the uninjured hemisphere and facilitating reorganization within the injured hemisphere, reinforcing its potential role as an adjunctive therapeutic agent in neuro-rehabilitation strategies. As an FDA-approved drug for mood stabilization, the potential benefit of Valproate as an adjuvant therapy alongside rehabilitation should be validated through robust clinical data. Retrospective studies on patients already receiving Valproate who later develop ischemic conditions could provide valuable insights into the drug’s neuromodulatory effects.

## 6 Acknowledgements

The authors wish to thank the National Institute of Mental Health and Neuro Sciences (NIMHANS) for providing infrastructure facilities to the candidate for carrying out the study. Sincere thanks to Dr. Ravi S. Muddashetty (Center for Brain Research, Bengaluru) for his insightful feedback and valuable guidance throughout the study. The authors would like to thank Mr. Gnan Kumar for his invaluable assistance, which contributed to the completion of this project. RPG contributed to the research design, experiments, analysis, and writing the original draft; DYN was involved in the study design and proofreading the manuscript; TRL was involved in Conceptualization, Study design, data analysis, review, and editing the manuscript

## 8. Data availability

All data analyzed during this study are included in this published article.

## 10. Ethical approval

Ethical approval was obtained from the Institutional Animal Ethics Committee (Approval ID: AEC/75/498/NP), which is under the guidelines of the Committee for the Purpose of Control and Supervision of Experiments on Animals (CPCSEA), Government of India.

## Bibliography

1. Wade DT, Langton-Hewer R, Wood VA, Skilbeck CE, Ismail HM. The hemiplegic arm after stroke: measurement and recovery. J Neurol Neurosurg Psychiatry. 1983 June 1;46(6):521–4.

2. Kwakkel G, Kollen BJ, Van Der Grond J, Prevo AJH. Probability of Regaining Dexterity in the Flaccid Upper Limb: Impact of Severity of Paresis and Time Since Onset in Acute Stroke. Stroke. 2003 Sept;34(9):2181–6.

3. Mutai H, Furukawa T, Nakanishi K, Hanihara T. Longitudinal functional changes, depression, and health-related quality of life among stroke survivors living at home after inpatient rehabilitation. Psychogeriatrics. 2016 May;16(3):185–90.

4. Post-stroke rehabilitation: assessment, referral, and patient management. U.S. Department of Health and Human Services Public Health Service. Agency for Health Care Policy and Research. Clin Pract Guidel Quick Ref Guide Clin. 1995 May;(16):i–iii, 1–32.

5. Gadidi V, Katz-Leurer M, Carmeli E, Bornstein NM. Long-Term Outcome Poststroke: Predictors of Activity Limitation and Participation Restriction. Arch Phys Med Rehabil. 2011 Nov;92(11):1802–8.

6. Mayo NE, Fellows LK, Scott SC, Cameron J, Wood-Dauphinee S. A Longitudinal View of Apathy and Its Impact After Stroke. Stroke. 2009 Oct;40(10):3299–307.

7. Lambert PA, Carraz G, Borselli S, Carbel S. [Neuropsychotropic action of a new anti-epileptic agent: depamide]. Ann Med Psychol (Paris). 1966 May;124(5):707–10.

8. Davis LL, Bartolucci A, Petty F. Divalproex in the treatment of bipolar depression: a placebo-controlled study. J Affect Disord. 2005 Apr;85(3):259–66.

9. Mathew NT, Saper JR, Silberstein SD, Rankin L, Markley HG, Solomon S, et al. Migraine Prophylaxis With Divalproex. Arch Neurol. 1995 Mar 1;52(3):281–6.

10. Evers S, Áfra J, Frese A, Goadsby PJ, Linde M, May A, et al. EFNS guideline on the drug treatment of migraine – revised report of an EFNS task force. Eur J Neurol. 2009 Sept;16(9):968–81.

11. Hao Y, Creson T, Zhang L, Li P, Du F, Yuan P, et al. Mood Stabilizer Valproate Promotes ERK Pathway-Dependent Cortical Neuronal Growth and Neurogenesis. J Neurosci. 2004 July 21;24(29):6590–9.

12. Shao L, Young LT, Wang JF. Chronic Treatment with Mood Stabilizers Lithium and Valproate Prevents Excitotoxicity by Inhibiting Oxidative Stress in Rat Cerebral Cortical Cells. Biol Psychiatry. 2005 Dec;58(11):879–84.

13. Christe W, Krämer G, Vigonius U, Pohlmann H, Steinhoff BJ, Brodie MJ, et al. A double-blind controlled clinical trial: oxcarbazepine versus sodium valproate in adults with newly diagnosed epilepsy. Epilepsy Res. 1997 Feb;26(3):451–60.

14. De Silva M, MacArdle B, McGowan M, Hughes E, Stewart J, Reynolds EH, et al. Randomised comparative monotherapy trial of phenobarbitone, phenytoin, carbamazepine, or sodium valproate for newly diagnosed childhood epilepsy. The Lancet. 1996 Mar;347(9003):709–13.

15. Chu BF, Karpenko MJ, Liu Z, Aimiuwu J, Villalona-Calero MA, Chan KK, et al. Phase I study of 5-aza-2′-deoxycytidine in combination with valproic acid in non-small-cell lung cancer. Cancer Chemother Pharmacol. 2013 Jan;71(1):115–21.

16. Takahashi S, Yoshida K, Paudel D, Morikawa T, Uehara O, Harada F, et al. Epigenetic agents, zebularine and valproic acid, inhibit the growth of the oral squamous cell carcinoma cell line HSC4 in vitro and in vivo. Discov Oncol. 2025 July 9;16(1):1293.

17. Brichta L. Valproic acid increases the SMN2 protein level: a well-known drug as a potential therapy for spinal muscular atrophy. Hum Mol Genet. 2003 Aug 5;12(19):2481–9.

18. Bach DR, Korn CW, Vunder J, Bantel A. Effect of valproate and pregabalin on human anxiety-like behaviour in a randomised controlled trial. Transl Psychiatry. 2018 Aug 16;8(1):157.

19. Aliyev NA, Aliyev ZN. Valproate (depakine-chrono) in the acute treatment of outpatients with generalized anxiety disorder without psychiatric comorbidity: Randomized, double-blind placebo-controlled study. Eur Psychiatry. 2008 Mar;23(2):109–14.

20. Smith LA, Cornelius VR, Azorin JM, Perugi G, Vieta E, Young AH, et al. Valproate for the treatment of acute bipolar depression: Systematic review and meta-analysis. J Affect Disord. 2010 Apr;122(1–2):1–9.

21. Keck PE, Taylor VE, Tugrul KC, McElroy SL, Bennett JA. Valproate treatment of panic disorder and lactate-induced panic attacks. Biol Psychiatry. 1993 Apr;33(7):542–6.

22. Adamou M, Puchalska S, Plummer W, Hale AS. Valproate in the treatment of PTSD: systematic review and meta analysis. Curr Med Res Opin. 2007 June;23(6):1285–91.

23. Neumann-Haefelin T, Staiger JF, Redecker C, Zilles K, Fritschy JM, Möhler H, et al. Immunohistochemical evidence for dysregulation of the GABAergic system ipsilateral to photochemically induced cortical infarcts in rats. Neuroscience. 1998 Aug;87(4):871–9.

24. Chen JY, Chu LW, Cheng KI, Hsieh SL, Juan YS, Wu BN. Valproate reduces neuroinflammation and neuronal death in a rat chronic constriction injury model. Sci Rep. 2018 Nov 7;8(1):16457.

25. Brookes RL, Crichton S, Wolfe CDA, Yi Q, Li L, Hankey GJ, et al. Sodium Valproate, a Histone Deacetylase Inhibitor, Is Associated With Reduced Stroke Risk After Previous Ischemic Stroke or Transient Ischemic Attack. Stroke. 2018 Jan;49(1):54–61.

26. Green AL, Zhan L, Eid A, Zarbl H, Guo GL, Richardson JR. Valproate increases dopamine transporter expression through histone acetylation and enhanced promoter binding of Nurr1. Neuropharmacology. 2017 Oct;125:189–96.

27. Kuruvilla A, Uretsky NJ. Effect of sodium valproate on motor function regulated by the activation of GABA receptors. Psychopharmacology (Berl). 1981 Jan;72(2):167–72.

28. Gibbons HM, Smith AM, Teoh HH, Bergin PM, Mee EW, Faull RLM, et al. Valproic acid induces microglial dysfunction, not apoptosis, in human glial cultures. Neurobiol Dis. 2011 Jan;41(1):96–103.

29. Qiu HM, Yang JX, Liu D, Fei HZ, Hu XY, Zhou QX. Antidepressive effect of sodium valproate involving suppression of corticotropin-releasing factor expression and elevation of BDNF expression in rats exposed to chronic unpredicted stress. NeuroReport. 2014 Mar 5;25(4):205–10.

30. Stout S. Effects of Sodium Valproate on Corticotropin-Releasing Factor Systems in Rat Brain. Neuropsychopharmacology. 2001 June;24(6):624–31.

31. Chen R, Xu M, Hogg RT, Li J, Little B, Gerard RD, et al. The Acetylase/Deacetylase Couple CREB-binding Protein/Sirtuin 1 Controls Hypoxia-inducible Factor 2 Signaling. J Biol Chem. 2012 Aug;287(36):30800–11.

32. Langley B, D’Annibale MA, Suh K, Ayoub I, Tolhurst A, Bastan B, et al. Pulse Inhibition of Histone Deacetylases Induces Complete Resistance to Oxidative Death in Cortical Neurons without Toxicity and Reveals a Role for Cytoplasmic p21 ^waf1/cip1^ in Cell Cycle-Independent Neuroprotection. J Neurosci. 2008 Jan 2;28(1):163–76.

33. Lin YH, Dong J, Tang Y, Ni HY, Zhang Y, Su P, et al. Opening a New Time Window for Treatment of Stroke by Targeting HDAC2. J Neurosci. 2017 July 12;37(28):6712–28.

34. Faraco G, Pancani T, Formentini L, Mascagni P, Fossati G, Leoni F, et al. Pharmacological Inhibition of Histone Deacetylases by Suberoylanilide Hydroxamic Acid Specifically Alters Gene Expression and Reduces Ischemic Injury in the Mouse Brain. Mol Pharmacol. 2006 Dec;70(6):1876–84.

35. Brookes RL, Crichton S, Wolfe CDA, Yi Q, Li L, Hankey GJ, et al. Sodium Valproate, a Histone Deacetylase Inhibitor, Is Associated With Reduced Stroke Risk After Previous Ischemic Stroke or Transient Ischemic Attack. Stroke. 2018 Jan;49(1):54–61.

36. Chen X, Wang H, Zhou M, Li X, Fang Z, Gao H, et al. Valproic Acid Attenuates Traumatic Brain Injury-Induced Inflammation in Vivo: Involvement of Autophagy and the Nrf2/ARE Signaling Pathway. Front Mol Neurosci. 2018 Apr 17;11:117.

37. Nesin SM, Sabitha KR, Gupta A, Laxmi TR. Constraint Induced Movement Therapy as a Rehabilitative Strategy for Ischemic Stroke—Linking Neural Plasticity with Restoration of Skilled Movements. J Stroke Cerebrovasc Dis. 2019 June;28(6):1640–53.

38. Liu P, Li C, Zhang B, Zhang Z, Gao B, Liu Y, et al. Constraint induced movement therapy promotes contralesional-oriented structural and bihemispheric functional neuroplasticity after stroke. Brain Res Bull. 2019 Aug;150:201–6.

39. Zhang A, Xing Y, Zheng J, Li C, Hua Y, Hu J, et al. Constraint-Induced Movement Therapy Modulates Neuron Recruitment and Neurotransmission Homeostasis of the Contralesional Cortex to Enhance Function Recovery after Ischemic Stroke. ACS Omega. 2024 May 14;9(19):21612–25.

40. Yue Y, Liu R, Cao Y, Wu Y, Zhang S, Li H, et al. New opinion on the subtypes of poststroke depression in Chinese stroke survivors. Neuropsychiatr Dis Treat. 2017 Mar;Volume 13:707–13.

41. Burton CAC, Murray J, Holmes J, Astin F, Greenwood D, Knapp P. Frequency of Anxiety after Stroke: A Systematic Review and Meta-Analysis of Observational Studies. Int J Stroke. 2013 Oct;8(7):545–59.

42. Li X, Wang X. Relationships between stroke, depression, generalized anxiety disorder and physical disability: some evidence from the Canadian Community Health Survey-Mental Health. Psychiatry Res. 2020 Aug;290:113074.

43. Kutlubaev MA, Hackett ML. Part II: Predictors of Depression after Stroke and Impact of Depression on Stroke Outcome: An Updated Systematic Review of Observational Studies. Int J Stroke. 2014 Dec;9(8):1026–36.

44. Totikov A, Boltzmann M, Schmidt SB, Rollnik JD. Influence of paroxysmal sympathetic hyperactivity (PSH) on the functional outcome of neurological early rehabilitation patients: a case control study. BMC Neurol. 2019 Dec;19(1):162.

45. Boyko M, Zlotnik A, Gruenbaum BF, Gruenbaum SE, Ohayon S, Goldsmith T, et al. An experimental model of focal ischemia using an internal carotid artery approach. J Neurosci Methods. 2010 Nov;193(2):246–53.

46. Morland C, Boldingh KA, Iversen EG, Hassel B. Valproate is Neuroprotective against Malonate Toxicity in Rat Striatum: An Association with Augmentation of High-Affinity Glutamate Uptake. J Cereb Blood Flow Metab. 2004 Nov;24(11):1226–34.

47. Sharma SS, Srinivas Bharath MM, Doreswamy Y, Laxmi TR. Effects of early life stress during stress hyporesponsive period (SHRP) on anxiety and curiosity in adolescent rats. Exp Brain Res. 2022 Apr;240(4):1127–38.

48. Sharma SS, Sasidharan A, Yoganarasimha D, Laxmi TR. Characterization of neuronal oscillations in the prelimbic cortex, nucleus accumbens and CA1 hippocampus during object retrieval task in rats predisposed to early life stress. Behav Brain Funct. 2024 Dec 18;20(1):34.

49. Garcia JH, Wagner S, Liu KF, Hu X Jiang. Neurological Deficit and Extent of Neuronal Necrosis Attributable to Middle Cerebral Artery Occlusion in Rats: Statistical Validation. Stroke. 1995 Apr;26(4):627–35.

50. Magno LA, Collodetti M, Tenza-Ferrer H, Romano-Silva M. Cylinder Test to Assess Sensory-Motor Function in a Mouse Model of Parkinson’s Disease. BIO-Protoc [Internet]. 2019 [cited 2024 Sept 26];9(16). Available from: https://bio-protocol.org/e3337

51. Sankaranarayani R, Nalini A, Rao Laxmi T, Raju TR. Altered neuronal activities in the motor cortex with impaired motor performance in adult rats observed after infusion of cerebrospinal fluid from amyotrophic lateral sclerosis patients. Behav Brain Res. 2010 Jan 5;206(1):109–19.

52. Zhang C, Zhang G, Liu D. Histone deacetylase inhibitors reactivate silenced transgene in vivo. Gene Ther. 2019 Apr;26(3–4):75–85.

53. DeBow SB, McKenna JE, Kolb B, Colbourne F. Immediate constraint-induced movement therapy causes local hyperthermia that exacerbates cerebral cortical injury in rats. Can J Physiol Pharmacol. 2004 Apr 1;82(4):231–7.

54. Ghosh B, Datta A, Gupta V, Sodnar B, Sarkar A, Singh U, et al. Simvastatin exerts neuroprotective effects post-stroke by ameliorating endoplasmic reticulum stress and regulating autophagy/apoptosis balance through pAMPK/LC3B/ LAMP2 axis. Exp Neurol. 2024 Nov;381:114940.

55. Ip Z, Rabiller G, He JW, Chavan S, Nishijima Y, Akamatsu Y, et al. Local field potentials identify features of cortico-hippocampal communication impacted by stroke and environmental enrichment therapy. J Neural Eng. 2021 Aug 1;18(4):0460a1.

56. Meneghetti N, Lassi M, Massa V, Micera S, Mazzoni A, Alia C, et al. Post-stroke spontaneous motor recovery in mice can be predicted from acute-phase local field potential using machine learning. APL Bioeng. 2025 June 1;9(2):026108.

57. Lawrence DG, Kuypers HGJM. THE FUNCTIONAL ORGANIZATION OF THE MOTOR SYSTEM IN THE MONKEY: I. THE EFFECTS OF BILATERAL PYRAMIDAL LESIONS. Brain. 1968;91(1):1–14.

58. Cortes JC, Goldsmith J, Harran MD, Xu J, Kim N, Schambra HM, et al. A Short and Distinct Time Window for Recovery of Arm Motor Control Early After Stroke Revealed With a Global Measure of Trajectory Kinematics. Neurorehabil Neural Repair. 2017 June;31(6):552–60.

59. Schambra HM, Xu J, Branscheidt M, Lindquist M, Uddin J, Steiner L, et al. Differential Poststroke Motor Recovery in an Arm Versus Hand Muscle in the Absence of Motor Evoked Potentials. Neurorehabil Neural Repair. 2019 July;33(7):568–80.

60. Cassidy JM, Wodeyar A, Wu J, Kaur K, Masuda AK, Srinivasan R, et al. Low-Frequency Oscillations Are a Biomarker of Injury and Recovery After Stroke. Stroke. 2020 May;51(5):1442–50.

61. Mizuhara H, Wang LQ, Kobayashi K, Yamaguchi Y. A long-range cortical network emerging with theta oscillation in a mental task: NeuroReport. 2004 June;15(8):1233–8.

62. Ondek K, Pevzner A, Tercovich K, Schedlbauer AM, Izadi A, Ekstrom AD, et al. Recovery of Theta Frequency Oscillations in Rats Following Lateral Fluid Percussion Corresponds With a Mild Cognitive Phenotype. Front Neurol. 2020 Dec 4;11:600171.

63. Rustamov N, Humphries J, Carter A, Leuthardt EC. Theta–gamma coupling as a cortical biomarker of brain–computer interface-mediated motor recovery in chronic stroke. Brain Commun. 2022 May 2;4(3):fcac136.

64. Perfetti B, Moisello C, Landsness EC, Kvint S, Lanzafame S, Onofrj M, et al. Modulation of Gamma and Theta Spectral Amplitude and Phase Synchronization Is Associated with the Development of Visuo-Motor Learning. J Neurosci. 2011 Oct 12;31(41):14810–9.

65. Brown P. Abnormal oscillatory synchronisation in the motor system leads to impaired movement. Curr Opin Neurobiol. 2007 Dec;17(6):656–64.

66. Tzagarakis C, Ince NF, Leuthold AC, Pellizzer G. Beta-Band Activity during Motor Planning Reflects Response Uncertainty. J Neurosci. 2010 Aug 25;30(34):11270–7.

